# Impacts of environmental factors on seed germination and seedling emergence of white clover (*Trifolium repens* L.)

**DOI:** 10.1101/2020.11.24.395681

**Authors:** Lei Chu, Yiping Gao, Lingling Chen, Patrick E. McCullough, David Jespersen, Suraj Sapkota, Xiao Li, Jialin Yu

**Author notes:** Lei Chu and Yiping Gao contributed equally to this work. Corresponding author’s (Jialin Yu).

## Abstract

White clover (*Trifolium repens* L.) is cultivated as a forage crop and planted in various landscapes for soil conservation. There are numerous reports of failed white clover stands each year. A good understanding of seed germination biology of white clover in relation to environmental factors is essential to achieve successful stand establishment. A series of experiments were conducted to investigate the impacts of light, temperature, planting depth, drought, and salt stress on seed germination and emergence of white clover. White clover is negatively photoblastic, and seed germination averaged 63 and 66% under light and complete dark conditions at 4 weeks after planting (WAP), respectively. Temperature affected seed germination speed and rate. At 1 WAP, seeds incubated at 15 to 25 °C demonstrated significantly higher germination rate than the low temperatures at 5 and 10 °C; however, the germination rate did not differ among the temperature treatments at 4 WAP. Results suggest that white clover germination decreases with increasing sowing depths and the seeds should be sown on the soil surface or shallowly buried at a depth ≤1 cm to achieve an optimal emergence. White clover seeds exhibited high sensitivity to drought and salinity stress. The osmotic potential and NaCl concentration required to inhibit 50% seed germination was −0.19 MPa and 62.4 mM, respectively. Overall, these findings provide quantifiable explanations for inconsistent establishment observed in field conditions. The findings obtained in this research can be used to develop effective planting strategies and support the successful establishment of white clover stands.

## 1 INTRODUCTION

White clover (*Trifolium repens* L.), a plant species in the leguminosae family, normally grows as a creeping, mulch-branched perennial (Frame and Newbould, 1986). White clover is found across a wide range of climates and distributed throughout the temperate and subtropical regions of the northern and southern hemispheres (Frame and Newbould, 1986; Uleberg et al., 2009). Due to its high crude protein content, white clover is widely cultivated as a legume forage crop (Baxter et al., 2019; Dazzo et al., 1978; Osoro et al., 2007). White clover is often included in perennial ryegrass (*Lolium perenne* L.) pastures (Black et al., 2006; Yu et al., 2008). This mixed system provides high-quality livestock feed (Osoro et al., 2007), while simultaneously improving soil fertility since legumes can fix atmospheric nitrogen with symbiotic bacteria (Elgersma et al., 2010).

In an intensively managed turfgrass system, such as a golf course, white clover is considered a problematic weed species. Nevertheless, for many low-maintenance turfgrass systems, such as airports, highway right-of-ways, residential lawns, and parks, the inclusion of white clover is recommended (McCurdy et al., 2013, 2014) and is a suggested strategy for improving the sustainability of low-maintenance turfgrasses (McCurdy et al., 2013, 2014). In China, white clover is often cultivated alone as a substitute for low maintenance turfgrasses (Ping et al., 2017) or intercropped into orchard and vineyard as a cover crop for the purpose of soil conservation (Wu et al., 2011; Song et al., 2006).

Unfortunately, there are numerous cases of inconsistent seedling emergence and stand establishments each year (Butler et al., 2014; Baxter et al., 2019; Laidlaw and Mcbride, 2010). The seed germination and seedling establishment of white clover might be highly correlated to soil and climatic conditions at time of planting. This has been demonstrated in many other legume species, such as chickpea (*Cicer arietinum* L.) (Ellis et al., 1986), red clover (*Trifolium pratense* L.) (Bukvic et al., 2010; Mandic et al. 2011), and yellow sweet clover (*Melilotus officinalis* L.) (Ghaderi-Far et al., 2010).

Seed germination and seedling emergence are the most important steps in a plant’s life cycle (Seneviratne et al., 2019; Yu et al. 2020). These stages are vulnerable to various abiotic stresses (Essemine et al., 2010; Koornneef et al., 2002). Therefore, understanding the germination characteristics of white clover sown under various environmental factors, such as seeding depth, drought, light, temperature, and salinity stress, may help develop effective sowing strategies. However, there is contradictory information on the effects of environmental factors on white clover germination and stand establishment reported in the literature. For example, Blaser and Killinger (1950) reported that white clover (cv. Louisiana) seed germination was poor under a continuous temperature of 22 °C or variable higher temperatures. However, in another study, Young et al. (1970) reported that sowing white clover seeds during warm weather early in the season promoted seed germination and successful stand establishment, while late rains and associated cold weather resulted in poor stand establishment. Monks et al. (2009) found that annual clover seed germination decreased after peaking at 20 °C, while perennial clover seed germination did not decrease until 35 °C.

There are many reports of failed white clover establishment each year (Butler et al. 2014; Caddel et al. 2004). White clover should be sown at optimal environmental conditions to avoid the soil and climatic factors that adversely affect seed germination and seedling emergence. Thus, there is a need to evaluate the impacts of various environmental factors on white clover seed germination and seedling emergence. The objective of this research was to examine the effects of light, temperature, seeding depth, drought, and salinity on white clover seed germination and seedling emergence.

## 2 MATERIALS AND METHODS

### 2.1 Seed germination test protocol

A series of experiments were conducted in spring and summer 2020 at the Nanjing Forestry University, Jiangsu Province, China (32.08 °N, 118.81 °E) to investigate the germination characteristics of white clover. An intermediate type of white clover was used in the study. The certified seeds (cv. Guizhou) were purchased from a commercial vendor (LvDing Seed Company, Suqian, Jiangsu, China). This white clover variety is presently being used for multiple purposes in China. It is planted with perennial ryegrass for grazing and is also planted in various landscapes for erosion control. The seeds were stored in dry conditions at 4 °C for 2 months until the initiation of experiments.

Unless otherwise specified, twenty-five white clover seeds were evenly placed in a Petri dish of 9 cm diameter containing two layers of filter paper (Whatman^®^ no. 1 filter paper, No. 6 Tianjing Nankai Rd, Tianjing, China) that was saturated with 5 ml of distilled water (pH = 7) or test solution. The Petri dishes were wrapped with parafilm and placed in controlled-environment growth chambers set for a 12-h photoperiod (light intensity 200 μmol m^-2^ s^-1^) with no diurnal temperature fluctuation and 70% relative humidity. Seeds were deemed to have germinated when protrusion of the radicle was visible.

### 2.2 Impact of temperature and light on seed germination

Experiments were conducted twice over time to evaluate the impact of temperature and light on white clover seed germination. The experiments were designed as a randomized complete block with four replications. The Petri dishes were maintained at a constant temperature of 5, 10, 15, 20, 25, or 30 °C under either 12-h photoperiod or complete dark. For the dark treatment, the Petri dishes were wrapped with aluminum foil to ensure no light penetration. Seed germination in darkness was assessed 4 weeks after planting (WAP).

### 2.3 Impact of burial depth on seedling emergence

Experiments were conducted twice over time in a controlled-environment growth chamber to evaluate the effect of seeding depth within the soil profile on seedling emergence. The experimental design was a randomized complete block with four replications. Twenty-five seeds were sown at 0 (soil surface), 1, 2, 3, 4, 6, 8, or 10 cm below the soil surface. The seeds were evenly scattered onto the surface of commercial potting soil (Dewoduofeiliao®, Dewoduo Fertilizer Co. Ltd., Heibei, China) and covered with field soil in square plastic pots (100 cm^2^ surface area by 15 cm height). The field soil was collected from a local farm and was classified as a Sandy loam (49.0% silt, 29.5% clay and 21.5% sand) with a pH 7.3 and 1% organic matter. The field soil was compressed by hand and leveled to standardize the burial depth. All pots were covered with polyethylene plastic film to prevent evaporation and to maintain soil moisture. Immediately after planting, the pots were placed in the environment-controlled growth chamber set for a 12-h photoperiod of 200 μmol m^−2^ s^−1^ and a constant temperature of 20 °C. Emerged seedlings were counted weekly for 4 weeks.

### 2.4 Impact of salinity stress on seed germination

Experiments were conducted twice over time to evaluate the impact of white clover seed germination at multiple levels of salt stress. The experimental design was a randomized block design with four replications. Salt solution treatments included 0, 10, 20, 40, 80, 160, and 320 mM NaCl. The Petri dishes were placed in the controlled-environment growth chamber set for a 12-h photoperiod with a constant temperature of 20 °C. Seed germination was counted weekly for 4 weeks or until no change in germination.

### 2.5 Impact of osmotic potential on seed germination

Experiments were carried out twice over time in environment-controlled growth chambers to evaluate white clover seed germination at multiple levels of osmotic potential. The experimental design was a randomized completed block with four replications. The osmotic potential treatments included 0, −0.0125 MPa (0.0199 g PEG ml^−1^ water), −0.025 MPa (0.0330 g PEG ml^−1^ water), – 0.05 MPa (0.0510 g PEG ml^−1^ water), −0.10 MPa (0.0778 g PEG ml^−1^ water), −0.15 MPa (0.0985 g PEG ml^−1^ water), −0.3 MPa (0.154 g PEG ml^−1^ water), −0.4 MPa (0.191 g PEG ml^−1^ water), −0.6 MPa (0.230 g PEG ml^−1^ water), −0.9 MPa (0.297 g PEG ml^−1^ water), −1.3 MPa (0.350 g PEG ml^−1^ water) (Michel, 1983). The Petri dishes were placed in growth chambers with a constant temperature of 20 °C and a 12-h photoperiod. Seed germination was counted weekly for 4 weeks.

### 2.6 Impact of pH on white clover seed germination

Experiments were conducted to evaluate the impact of pH on white clover seed germination. Six pH buffered solutions were prepared using a method described by Chen et al. (2009). The seeds were exposed to aqueous solutions of pH 4 (10^−4^ mol L^−1^ HCl), pH 5 (10^−5^ mol L^−1^ HCl), pH 6 (10^−6^ mol L^−1^ HCl), pH 8 (10^−6^ mol L^−1^ NaOH), pH 9 (10^−5^ mol L^−1^ NaOH), pH 10 (10^−4^ mol L^−1^ NaOH), and deionized water (pH 7). The Petri dishes were placed in growth chambers set for a constant temperature of 20 °C and a 12-h photoperiod. Seed germination was counted 4 WAP.

### 2.7 Statistical analyses

Data were subjected to analysis of variance (ANOVA) with the MIXED procedure in SAS (v. 9.4, SAS Institute Inc., Cary, NC). The experimental run and blocks were considered as random factors, while treatments were considered as fixed factors. Where data measurements were conducted over time, data were subjected to repeated measures ANOVA. Homogeneity of equal variance and normality were checked prior to analysis. Data were log-transformed when necessary but back-transformed data are presented. One-way ANOVA was performed to analyze the impacts of temperature on seed germination, while two-way ANOVA was performed to analyze the impacts of temperature and light on seed germination. Treatment means were compared with Tukey’s adjustment means comparison at α = 0.05.

The seed germination values from osmotic potentials, salt stress, or seedling emergence data from burial depths were regressed with a functional three-parameter sigmoid model. The model fitted was:

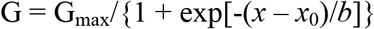

where G is the total germination or emergence (%) at the osmotic potential (MPa), NaCl concentration (mM), or burial depth (cm) *x*, G_max_ is the maximum germination or emergence (%), *x*_0_ is the osmotic potential, NaCl concentration, or burial depth needed for inhibiting 50% maximum germination or emergence, and *b* represents the slope (Ritz et al., 2015).

## 3 RESULTS AND DISCUSSION

### 3.1 Impact of temperature and light and on seed germination

The repeated measures analysis indicated that the measurement timing by temperature interaction was significant (*P* < 0.05) and therefore measurement timings are presented separately (Table 1). In the presence of light, the seed germination rate at 1 WAP was 4 and 48% at 5 and 10 °C, respectively, while the germination rate was ≥61% at the temperatures ranging from 15 to 25 °C. The germination rate at 30 °C was 60% and was higher than 5 °C but did not differ from temperatures ranging from 10 to 25 °C. The germination rate was 5 and 29% at 2 and 3 WAP at 5 °C, respectively, and was lower compared to higher temperatures ranging from 10 to 30 °C.

**TABLE 1.** Impact of light and temperature on white clover seed germination in environment-controlled growth chambers.

| Temperature (°C) | Germination |  |  |  |  |
| --- | --- | --- | --- | --- | --- |
|  | Light |  |  |  | Dark |
|  | 1 WAP | 2 WAP | 3 WAP | 4 WAP | 4 WAP |
|  | -----%----- |  |  |  |  |
| 5 | 4 ± 1.4 c | 5 ± 2.3 b | 29 ± 6.8 b | 62 ± 6.8 | 60 ± 4.8 |
| 10 | 48 ± 3.2 b | 54 ± 3.9 a | 59 ± 3.8 a | 61 ± 3.8 | 71 ± 4.2 |
| 15 | 67 ± 3.9 a | 67 ± 3.9 a | 69 ± 4.1 a | 69 ± 4.1 | 64 ± 4.1 |
| 20 | 62 ± 2.3 a | 62 ± 2.3 a | 62 ± 2.6 a | 62 ± 2.6 | 64 ± 2.6 |
| 25 | 61 ± 3.5 a | 62 ± 3.6 a | 62 ± 3.6 a | 62 ± 3.6 | 65 ± 3.6 |
| 30 | 60 ± 3.7 ab | 60 ± 3.7 a | 60 ± 4.9 a | 60 ± 4.9 | 69 ± 4.9 |
| *P*-value | <0.0001 | <0.0001 | <0.0001 | 0.6745 | 0.3068 |

<sup>a</sup>Treatment means within the same column were not significantly different according to one-way Tukey's adjustment means comparison at the 0.05 significance level. Data represent the treatment means ± standard errors (n = 8). Results were pooled over two experimental runs.
<sup>b</sup>Abbreviation: WAP = weeks after planting.

At 4 WAP, the impacts of temperature and light condition and their interaction on seed germination were not significant (*P* > 0.05) (data not shown). The seed germination rate in the presence of light or complete darkness did not differ among the temperatures (Table 1). The germination rate averaged 63 and 66% under light and complete dark, respectively. The germination occurred under both light and dark conditions, suggesting that white clover is negatively photoblastic and thus light is not required for germination.

These results support the idea that sowing white clover at an appropriate temperature range is critical to achieve rapid germination (Baxter et al. 2019). Planting seeds at ≥15 °C is desirable and should result in early seed germination and seedling establishment, provided that other environmental factors are suitable. However, this finding does not necessarily imply white clover seeds should be sown in warm weather. In previous research, Blaser and Killinger (1950) noted that the prevailing high temperatures resulted in low or even absent white clover populations in a pasture in Florida. This is likely due to the fact that white clover can achieve a high level of leaf expansion at low temperature and light intensity (Frame and Newbould, 1986) and grows best during cool, moist weather (Anonymous, 2020).

The results suggest that timing of planting is critical to accomplish rapid germination and successful stand establishment. The midsummer in parts of China is often associated with extreme high temperatures and soil moisture deficits. The late-fall sowing prior to the onset of low winter temperatures, especially in northern regions of China, may result in insufficient time for adequate establishment. Therefore, spring is the best time for sowing white clover. However, caution should to be taken when sowing too early in spring because of the slow germination at low temperatures.

Our findings suggest that the optimal temperature range for germination is 15 to 25 °C. In previous research, Kendall and Stringer (1985) noted that the optimum germination temperature for annual clovers was 15 °C. However, Butler et al. (2014) noted that the optimum germination for perennial legume clovers was around 20 °C. In a more recent study, Baxter et al. (2019) found that the optimal germination temperature range for white clover (cv. Neches) was 10.9 to 17.2 °C. Seeds of clover varieties originating in northern locations generally exhibited high tolerance to low temperatures compared to varieties naturally selected in southern locations (Hoveland and Elkins, 1965). It should be mentioned that the clover variety examined in our study may exhibit dissimilar germination patterns compared to other varieties in response to temperature, which warrants further investigation.

### 3.2 Impact of sowing depth on seedling emergence

A sigmoidal response was observed with seedlings emergence of white clover as sowing depth increased (Figure 1). The highest emergence was consistently observed when seeds were sown on the soil surface but decreased when seeds were sown at deeper depths. We also noticed faster seedling emergence when seeds were planted on the soil surface or buried at shallower depths compared to deeper depths. At 1 WAP, emergence was 20% when the seeds were sown on the soil surface, while no seedlings emerged from a depth of 1 cm or deeper. At 2 WAP, emergence was 63% when the seeds were sown on the soil surface but never exceeded 50% at 1 cm depth or deeper. The emergence stabilized at 3 WAP because emerged seedlings across burial depths were identical between 3 and 4 WAP. At 3 and 4 WAP, the emergence was 77% when the seeds were planted on the soil surface but declined to 55, 59, and 40% when the seeds were sown at the 1, 2, and 3 cm depths, respectively. Emergence never exceeded 23% for the seeds buried at a depth of 4 cm or greater.

**FIGURE 1.**
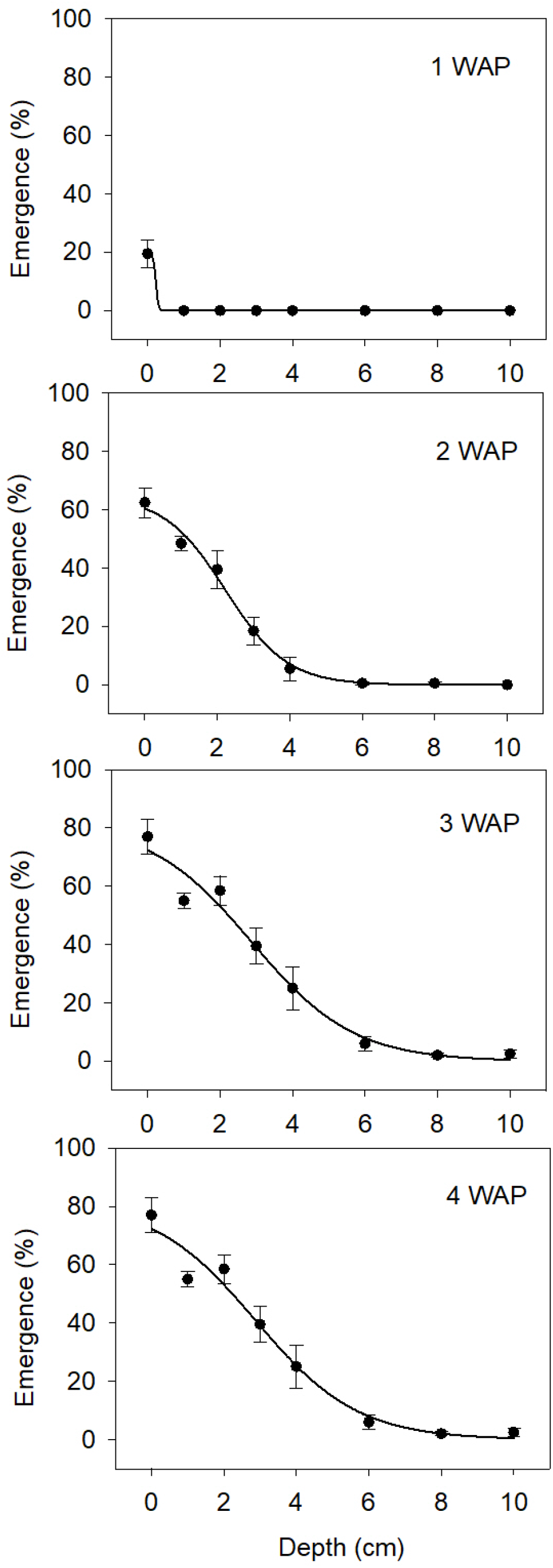
White clover seed germination over time at various burial depths. Data were regressed with the following equation: G = G_max_/{1 + exp[-(x – x_0_)/*b*]}. Equation for 1 WAP was not available, *P* > 0.05 did not achieve a significant nonlinear relationship. Equation for 2 WAP was G = 64.89/{1 + exp[-(x – 2.22)/−0.85]}, standard errors for G_max_, x_0_, *b* measured 6.51, 0.26, and 0.19, respectively, R^2^ = 0.84, and *P* < 0.0001. Equation for 3 WAP was G = 78.84/{1 + exp[-(x – 2.86)/−1.33]}, standard errors for G_max_, x_0_, *b* measured 10.62, 0.49, and 0.34, respectively, R^2^ = 0.80, and *P* < 0.0001. Equation for 4 WAP was G = 81.12/{1 + exp[-(x – 2.85)/−1.35]}, standard errors for G_max_, x_0_, *b* measured 10.76, 0.49, and 0.34, respectively, R^2^ = 0.81, and *P* < 0.0001. Results were pooled over two experimental runs. Vertical bars represent standard errors (n = 8). Abbreviation: WAP = weeks after planting.

Our results clearly indicate that white clover seeds should be sown on the soil surface or at ≤1 cm depth to maximize seedling emergence. Sowing white clover too deep can cause poor seedling establishment. This is probably because white clover is a small-seeded species, with limited energy reserves, and thus insufficient seed reserves failed to support seedling emergence from deeper soil depths. This has been demonstrated in many small-seeded plant species such as black medic (*Medicago lupulina* L.), common beggar’s-tick (*Bidens alba* L.), cobbler’s pegs (*Bidens Pilosa* L.), and sweet clover (*Meliotus officinalis* L.) (Chauhan et al., 2019; Ghaderi-Far et al., 2010; Sharpe and Boyd, 2019; Yu et al., 2020). The recommended white clover sowing depth is 0.3–0.6 cm (USDA-NRCS Plant Guide, 2020). This recommendation may have taken into account the fact that the seeds that are sown on the soil surface prone to environmentally facilitated death, including insect predation. In addition, deceases in seedling emergence due to increasing burial depth is also related to temperature and soil moisture (Bliss and Smith, 2010; Stoller, 1978; Sharpe and Boyd, 2019; Yu et al., 2020). Further analysis needed to evaluate possible interactions between temperature, soil moisture, and burial depth on white clover emergence.

### 3.3 Impact of salinity stress on germination

A sigmoidal response was observed in the germination of white clover seeds with increases in NaCl concentrations ranging from 0 to 320 mM (Figure 2). The highest germination was observed when the seeds were incubated in the solutions of non-stress distilled water. While germination was completely inhibited at ≥160 mM NaCl, 15% of seeds germinated at 80 mM NaCl. The NaCl concentration required to inhibit 50% seed germination was 62 mM.

**FIGURE 2.**
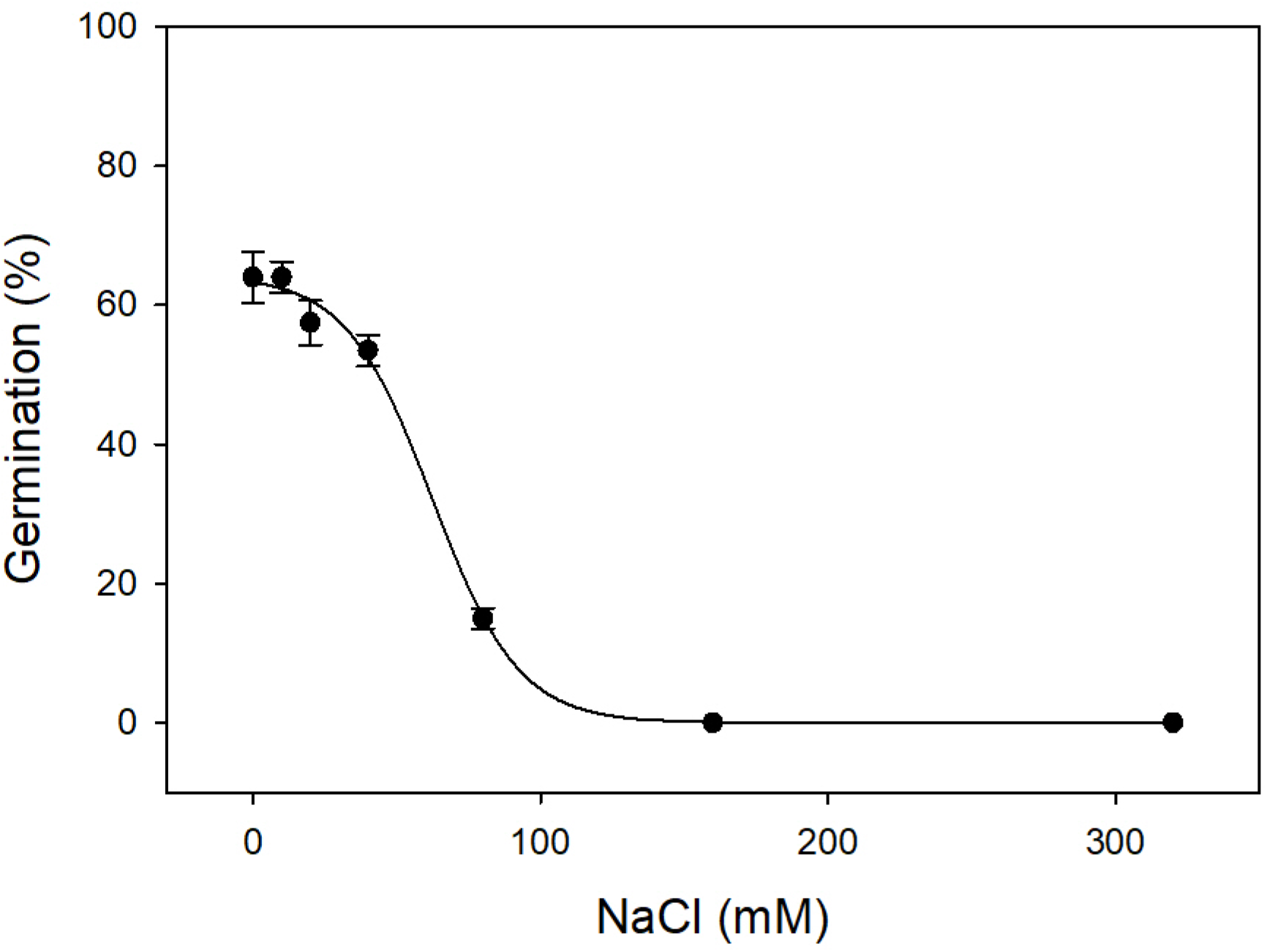
White clover seed germination over time at various NaCl concentrations. Data were regressed with the following equation: G = G_max_/{1 + exp[-(x – x_0_)/*b*]}. Equation was G = 64.29/{1 + exp[-(x – 62.40)/−15.02]}, standard errors for G_max_, x_0_, *b* measured 1.76, 2.46, and 1.76, respectively, R^2^ = 0.96, and *P* < 0.0001. Results were pooled over two experimental runs. Vertical bars represent standard errors (n = 8). Abbreviation: WAP = weeks after planting.

**FIGURE 3.**
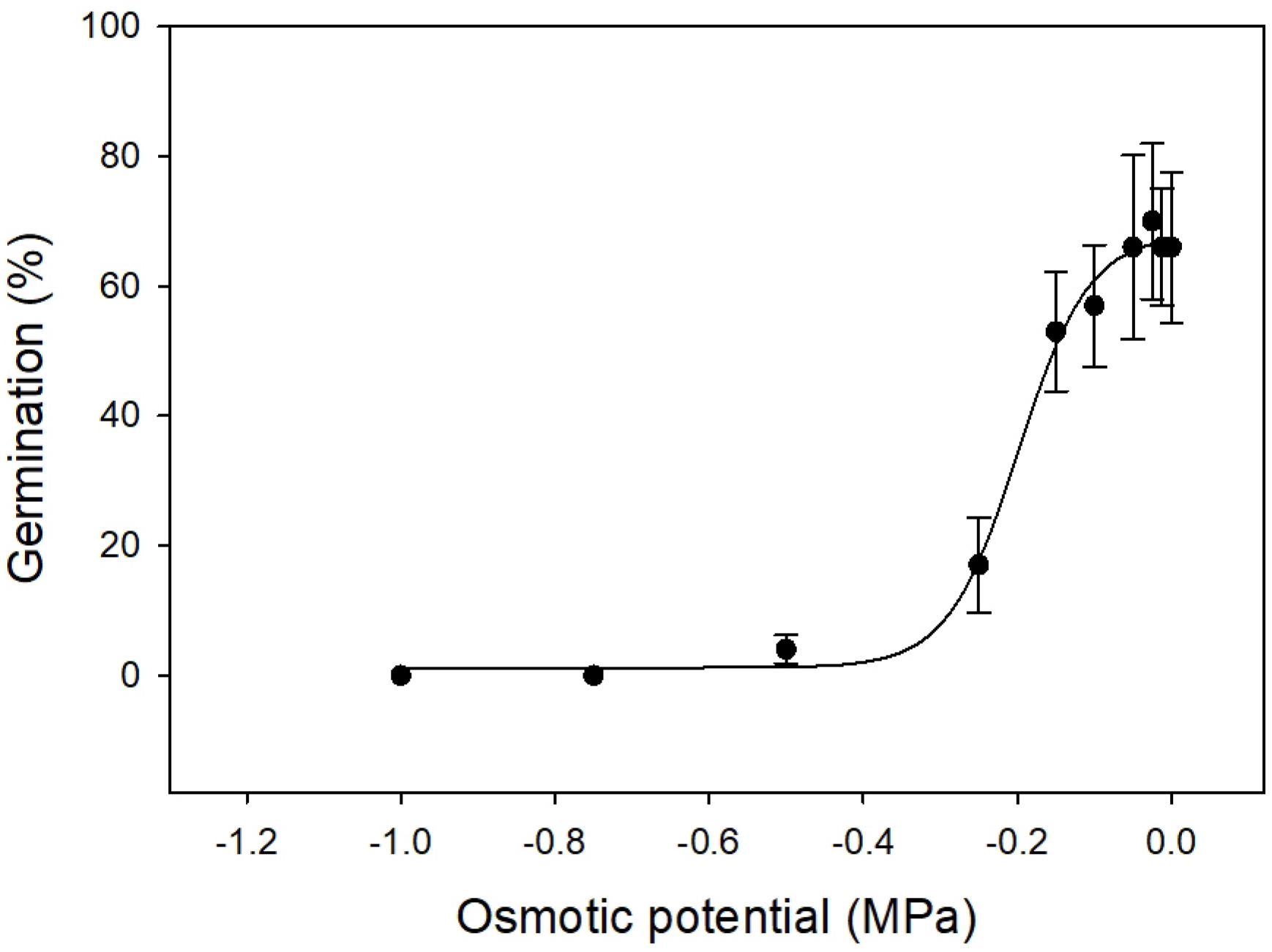
White clover seed germination over time at various osmotic potentials (MPa). Data were regressed with the following equation: G = G_max_/{1 + exp[-(x – x_0_)/*b*]}. Equation was G = 67.74/{1 + exp[-(x – −0.1997)/−0.0469]}, standard errors for G_max_, x_0_, *b* measured 4.56, 0.0174, and 0.0144, respectively, R^2^ = 0.79, and *P* < 0.0001. Results were pooled over two experimental runs. Vertical bars represent standard errors (n = 8). Abbreviation: WAP = weeks after planting.

These results confirmed that seed germination of white clover is susceptible to salinity stress. Seed germination and seedling establishment are the most vulnerable plant growth stages to salinity stress (Munns and Tester, 2008; Masood et al., 2018). Salinity stress can cause a series of harmful events including ionic imbalances, nutritional imbalances, and excessive accumulation of reactive oxygen species (Parihar et al., 2015; Yadav et al., 2011). Moreover, the overaccumulation of Cl^−^ and Na^+^ can induce osmotic stress and reduce water uptake, resulting in physiological drought (Cheng et al., 2018; Munns and Tester, 2008). All these responses can contribute to the deleterious effects on plant growth and development. In other legume species, salinity stress substantially reduced seed germination and seedling emergence (Chatterjee, 2018; Ghaderi-Far et al., 2010; Yilmaz and Kulaz, 2019; Sehrawat et al., 2019).

In previous investigations, Guo et al. (2013) reported that seed priming with silicon effectively alleviated sodium toxicity in white clover by enhancing the selective transport capacity for K+ over Na+, resulting in reduced Na+ uptake and increased K+ uptake by roots. Cheng et al. (2018) reported that seed priming with γ-aminobutyric acid effectively mitigated salt-induced stress and improved seed germination of white clover (cv. Haifa). Further research is needed to evaluate the effect of seed priming on seed germination and seedling emergence for the present white clover variety.

### 3.4 Impact of osmotic potential on white clover germination

A sigmoidal response was noted in the germination of white clover seeds with reductions in osmotic potential from 0 to −1 MPa. The low osmotic potentials ranging from 0 to −0.05 MPa did not affect the germination rate. However, the germination sharply decreased from 57 to 17% as osmotic potential dropped from −0.01 MPa to −0.025 MPa. The germination rate was only 4% when the seeds were incubated at −0.5 MPa. No germination occurred at an osmotic potential of − 0.75 MPa. The osmotic potential required to inhibit 50% germination was −0.19 MPa.

In other legume species, black medic (*Medicago lupulina* L.) exhibited a similar germination response to varying osmotic potentials (Sharpe and Boyd, 2019). Osmotic stress can also unfavorably influence the legume plant seedling establishment by reducing net photosynthesis and increasing the production of reactive oxygen species (Farooq et al., 2016). Osmotic stress negatively impacted the metabolic pathways of alfalfa (Naya et al., 2007) and cowpea (*Vigna unguiculate* L. Walp.) (Figueiredo et al., 2007) seedlings by suppressing several enzymes, including glutamate synthase, phosphoenolpyruvate carboxylase, and sucrose synthase, involved in nitrogen fixation. Overall, the findings in the present study suggest that white clover seed germination is very sensitive to drought. Drought could result in poor germination and high mortality rate. Therefore, white clover seeds should only be planted on moist soil or shortly prior to the rainy weather.

### 3.5 Impact of pH on white clover germination

White clover seed germination did not differ among the pH treatments (*P* > 0.05) (Figure 4). Among the tested pH solutions, at least 68% white clover seeds were germinated. This result suggests that white clover seeds can germinate successfully over a wide range of pH conditions. It is well known that soil pH can affect plant seed germination and emergence (Boyd et al. and Hughes 2011; Caddel et al. 2004). Nevertheless, soil pH may decrease with time due to acidic rainfall, organic matter decay, and nitrogen fertilization (Sparks et al. 1996). In previous research, growth of white clover in sand culture under symbiotic conditions at pH 4 was nil but increased linearly from pH 4 to 6 (Andrew 1976; Snaydon (1962).

**FIGURE 4.**
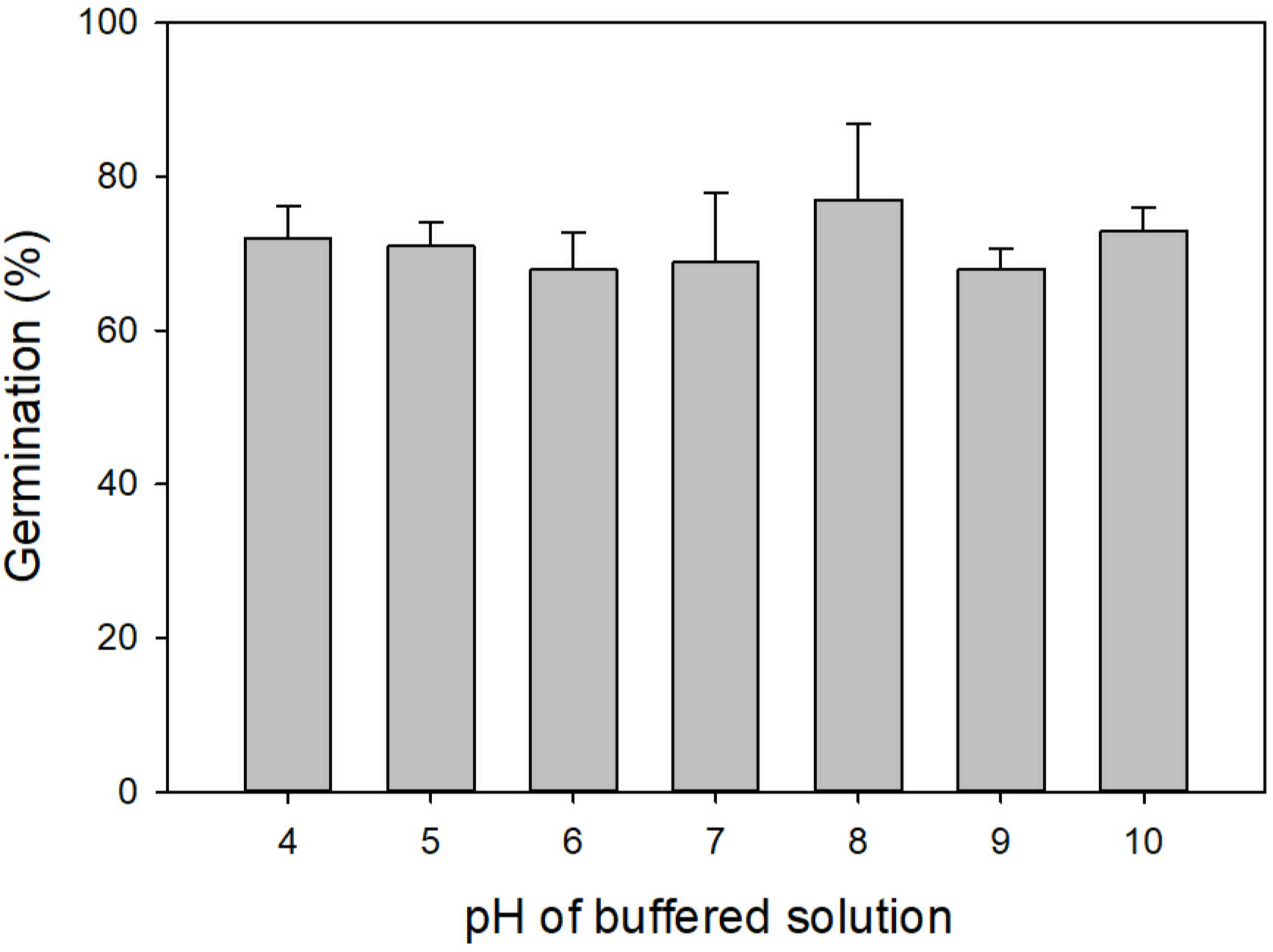
Impact of buffered pH solutions on white clover seed germination incubated in growth chambers set for a constant temperature of 20 °C and a 12-h photoperiod with a light intensity of 200 μmol m^−2^ s^−1^ and 70% relative humidity. Results were pooled over two experimental runs. Vertical bars represent standard errors (n = 8).

## 5 Implications from these findings

This research demonstrated that temperature, sowing depth, drought, and salinity stress can significantly affect the successful establishment of white clover. Regarding possible strategies in achieving successful white clover establishment, the following issues should be noted. Temperature is one of the main factors affecting the speed and rate of white clover germination. Increased sowing depth can reduce seedling emergence. White clover should be sown on the soil surface or shallowly incorporated into the soil profile at a temperature ranging from 15 to 25 °C for achieving optimum emergence time and rate. Because of the high sensitivity to salinity stress, white clover should not be sown onto the salt-affected soils or irrigated with salt-loaded water. White clover seed germination is very sensitive to drought stress. A moist soil is critical following planting to achieve a successful establishment. Seed germination of white clover over a broad pH range suggests that soil pH is not a limiting factor for germination. The results obtained in the present study can be employed to develop appropriate strategies and support the successful establishment of white clover.

## Acknowledgments

The authors would like to thank Lu Yang for technical assistance. This research received no specific grant from any funding agency in the commercial, public, or not-for-profit organizations.

## Data Availability

No experimental data described.

